# Efficient inhibition of SARS-CoV-2 strains by a novel ACE2-IgG4-Fc fusion protein with a stabilized hinge region

**DOI:** 10.1101/2020.12.06.413443

**Authors:** Hristo L. Svilenov, Julia Sacherl, Alwin Reiter, Lisa Wolff, Cho-Chin Chen, Marcel Stern, Frank-Peter Wachs, Nicole Simonavicius, Susanne Pippig, Florian Wolschin, Johannes Buchner, Carsten Brockmeyer, Ulrike Protzer

## Abstract

The novel severe acute respiratory syndrome (SARS)-like coronavirus (SARS-CoV-2) enters its host cells after binding to the angiotensin-converting enzyme 2 (ACE2) *via* its spike glycoprotein. This interaction is critical for virus entry and virus-host membrane fusion. Soluble ACE2 ectodomains bind and neutralize the virus but the short *in vivo* half-lives of soluble ACE2 limits its therapeutic use. Fusion of the fragment crystallizable (Fc) part of human immunoglobulin G (IgG) to the ACE2 ectodomain can prolong the *in vivo* half-life but bears the risk of unwanted Fc-receptor activation and antibody-dependent disease enhancement. Here, we describe optimized ACE2-Fc fusion constructs that avoid Fc-receptor binding by using IgG4-Fc as a fusion partner. The engineered ACE2-IgG4-Fc fusion proteins described herein exhibit promising pharmaceutical properties and a broad antiviral activity at single-digit nanomolar concentration. In addition, they allow to maintain the beneficial enzymatic activity of ACE2 and thus are very promising candidate antivirals broadly acting against coronaviruses.

## Introduction

Of the seven known human pathogenic coronavirus (hCoV) strains, three (SARS-CoV, SARS-CoV-2, hCoV-NL63) use the angiotensin-converting enzyme 2 (ACE2) as a receptor for entry into the human host cell [1–3]. Neuropilin-1 has recently been identified as another protein supporting SARS-CoV-2 infection potentially handing the virus over to ACE2 [4]. ACE2 is located in the plasma membrane of respiratory epithelial cells [3]. In addition to the respiratory tract, high expression of ACE2 is also found in several other tissues, including intestine, testes, liver, kidney, brain, and the cardiovascular system [5–7].

ACE2 is a 805 amino acid type-I transmembrane protein consisting of an extracellular, a transmembrane and a cytosolic domain [8]. The extracellular domain is a zinc metalloprotease, which enzymatically functions as a carboxypeptidase [9, 10] suppressing the renin-angiotensin system (RAS) by cleaving angiotensin I to the inactive angiotensin 1-9 and even more efficiently cleaving angiotensin II to angiotensin 1-7 [11, 12]. Angiotensin 1-7 and its downstream peptides exert a broad spectrum of cell and tissue protection [13], it lowers diastolic blood pressure, has anti-inflammatory, anti-proliferative and anti-fibrotic effects and thereby protects the lung, heart, kidney and other organs from injury [5–7]. The potential contribution of angiotensin II to the COVID-19 pathophysiology has been indicated by reports that angiotensin II levels in plasma samples from COVID-19 patients were markedly elevated and correlated with viral load and severity of disease [14, 15].

ACE2-Fc fusion proteins composed of the human immunoglobulin G (IgG) fragment crystallizable (Fc) part fused to the extracellular domain of ACE2 have been suggested as a high-priority treatment option for COVID-19 [16, 17]. In addition to neutralizing SARS-CoV and SARS-CoV-2, the ACE2 enzymatic activity affecting the renin-angiotensin system provides a second mode of action, potentially alleviating the pathophysiology of acute respiratory distress syndrome (ARDS). Therapeutic use of a soluble human recombinant ACE2 dimer (APN01) with a half-life of 10 hours is currently investigated in patients with COVID-19 [18–20]. Strong *in vitro* SARS-CoV-2 neutralizing activity has been described for several sequence variants of ACE2-IgG1-Fc fusion proteins [21–26]. A version of an ACE2-IgG1-Fc (HLX71) is in a Phase I clinical study with healthy volunteers [27].

A concern arising from the experience with vaccines and neutralizing antibodies is disease enhancement by Fc effector functions such as complement dependent cytotoxicity (CDC) and antibody dependent cytotoxicity (ADCC) [28]. Moreover, Fc receptor gamma III (CD16) binding, which for example has been shown for neutralizing antibodies in Middle East respiratory syndrome (MERS), has led to infection of CD16 positive cells [29, 30]. It is well known that IgG1-Fc strongly binds to CD16 and has pronounced CDC and ADCC activity, whereas in contrast for IgG4-Fc such Fc-related effector functions are minimal [31]. For this reason, the IgG4-Fc fragment would be a preferred fusion partner for ACE2. However, it is well known that naturally occurring IgG4 antibodies are less stable than IgG1 variants due to the formation of half antibodies, which limits their use in pharmaceutical preparations [32–35].

Therefore, we have chosen the immunoglobulin Fc region of an IgG4/kappa isotype with an S228P sequence alteration in the hinge region to generate a stable ACE2-IgG4-Fc fusion protein [33]. For the ACE2 domain two different truncations, Q18-G732 and Q18-S740, respectively, were used for the fusion. In addition, in alternative constructs, point mutations were introduced in the ACE2 domain to abrogate its enzymatic activity.

Here we report the biophysical and functional characteristic of these ACE2-IgG4-Fc fusion proteins and demonstrate that they efficiently target and entirely neutralize different strains of SARS-CoV-2 circulating in Europe in January and April 2020 as well as the 2003 SARS-CoV with strain-specific high affinities.

## Results

### Expression and purification of ACE2-Fc proteins

ACE2-IgG4-Fc and ACE2-IgG1-Fc fusion proteins were designed based on crystal and EM structures of the ACE2 extracellular domain, the SARS-CoV-2 spike (S) protein and its receptor binding domain (RBD), as well as the IgG4-Fc and IgG1-Fc domains [3, 36–38] (**Figure 1a and b**). Details of the ACE2 sequences fused to the Fc fragments of IgG4 and IgG1 are shown in **Table 1.**

**Figure 1.**
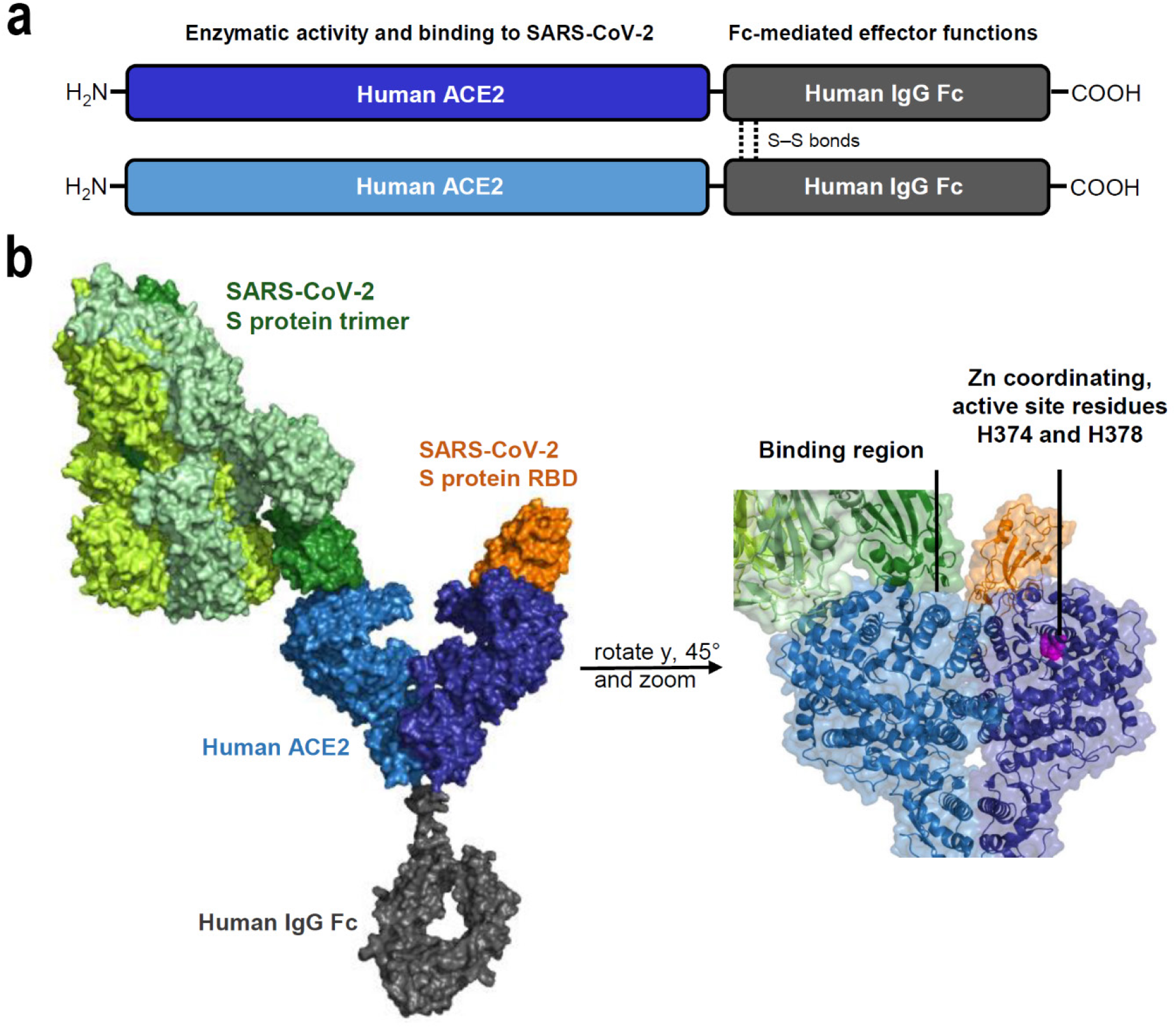
Structural elements in the ACE2-Fc constructs. **a** Schematic depiction of the main parts in an ACE2-Fc molecule and their functional properties. **b** Design of the ACE2-Fc fusion protein; ACE2 parts in light and dark blue, IgG-Fc part in gray, spike (S) protein trimer in light green and the receptor binding domain (RBD) located at the tip of each spike protein in orange and dark green. The binding region as well as active site residues H374 and H378 important for the enzymatic activity of ACE2 are highlighted. Structures of the following Protein Data Bank (PBD) identifiers were used for modeling: 6M17, 6M0J, 6VSB, 5DK3.

**Table 1.** Structural properties of the ACE2-Fc constructs

| Construct | ACE2 sequence (active site mutation) | IgG isotype (hinge region modification) |
| --- | --- | --- |
| 1 | Q18-G732 (no) | IgG4 (S228P) |
| 2 | Q18-S740 (no) | IgG4 (S228P) |
| 3 | Q18-G732 (H374N, H378N) | IgG4 (S228P) |
| 4 | Q18-S740 (H374N, H378N) | IgG4 (S228P) |
| 5 | Q18-G732 (no) | IgG1 (truncated) |
| 6 | Q18-S740 (no) | IgG1 (truncated) |
| 7 | Q18-G732 (H374N, H378N) | IgG1 (truncated) |
| 8 | Q18-S740 (H374N, H378N) | IgG1 (truncated) |

The expression yields of all fusion proteins were in a similar range, although slightly higher for Q18-G732 ACE2-Fc fusion proteins. Purity of the ACE2-Fc fusion proteins under investigation ranged between 95% and 98%, as evident from capillary electrophoresis sodium dodecyl sulfate (CE-SDS) and size exclusion chromatography (SEC) analysis. High molecular weight fractions (HMWS) measured after purification were slightly lower for Q18-G732 ACE2-Fc fusion proteins (**Supplementary Table 1**). Peptide mapping confirmed the presence of the modifications in the ACE2-Fc fusion proteins.

### Variations in the ACE2-Fc sequences have a minor effect on the basic structural properties

To assign secondary structure to the ACE2-Fc fusion construct, protein circular dichroism (CD) spectroscopy was performed. The Far-UV (190-250 nm) CD spectra of the fusion proteins could be superimposed indicating that the secondary structures are preserved among all constructs, regardless of the sequence variations **(Figure 2a left)**. The same held true for the Near-UV (250- 300 nm) CD spectra, which indicated that the overall tertiary structure is also highly similar in all ACE2-Fc proteins investigated **(Figure 2a right)**.

**Figure 2.**
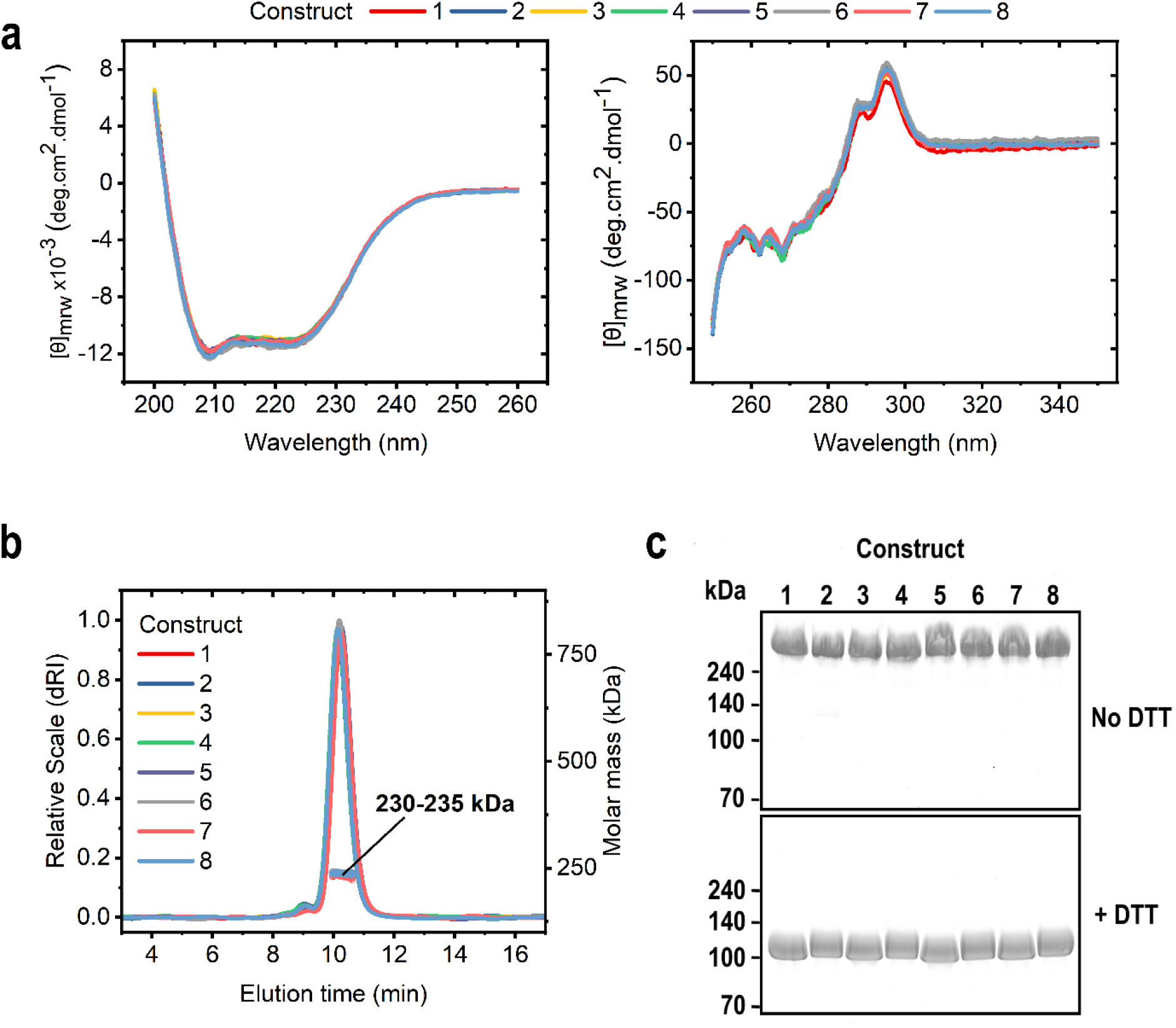
Structural characterization of the ACE2-Fc proteins. **a** Far-UV CD spectra (left) and Near-UV CD spectra (right) of ACE2-Fc constructs indicating that all proteins exhibit same secondary and tertiary structures. **b** Chromatograms and molecular mass from size-exclusion chromatography coupled to multi-angle light scattering (SEC-MALS) indicating that the ACE2-Fc molecules form homodimers. **c** Non-reducing (top) and reducing (bottom) sodium dodecyl sulfate polyacrylamide gel electrophoresis (SDS-PAGE) analysis showing that intermolecular disulfide bonds in the homodimers are formed.

Size-exclusion chromatography coupled to multi-angle light scattering (SEC-MALS) was used to investigate the oligomeric state of the fusion proteins. All ACE2-Fc fusion molecules tested exhibit similar elution times and molecular masses of 230 to 235 kDa (**Figure 2b)**. These values agreed well with the theoretical mass of a homodimer, which is 216-218 kDa based on calculations from the primary sequence. The slightly higher molecular mass measured in solution was due to the glycosylation of the proteins.

Reducing and non-reducing sodium dodecyl sulfate polyacrylamide gel electrophoresis (SDS-PAGE) **(Figure 2c)** revealed that all ACE2-Fc fusion proteins separated under reducing conditions with dithiothreitol (DTT) run at about 110-120 kDa, which correlated to the theoretical mass of a monomeric ACE2-Fc molecule. The non-reduced ACE2-Fc fusion proteins exhibited a much higher apparent molecular weight, indicating that the intermolecular disulfide bonds in the ACE2-Fc homodimers are formed, which agrees with results from CE-SDS under reducing and nonreducing conditions.

### H374N and H378N mutations abolish enzymatic activity of ACE2-Fc constructs

The *in vitro* assay for the catalytic activity of ACE2 is based on the cleavage of a synthetic peptidyl-4-methylcoumaryl-7-amide (MCA) and the release of the free MCA fluorophore that has an increased fluorescence intensity at 420 nm (excitation at 320 nm) compared to the peptidyl-MCA. The amount of released MCA was calculated from the slope of the increase in fluorescence intensity and a standard curve with known MCA concentrations. All constructs with wildtype (WT) ACE2 cleave the same amount of MCA during 30 min of incubation, while the constructs with mutations in the active ACE2 site lost the enzymatic activity **(Figure 3a and b)**.

**Figure 3.**
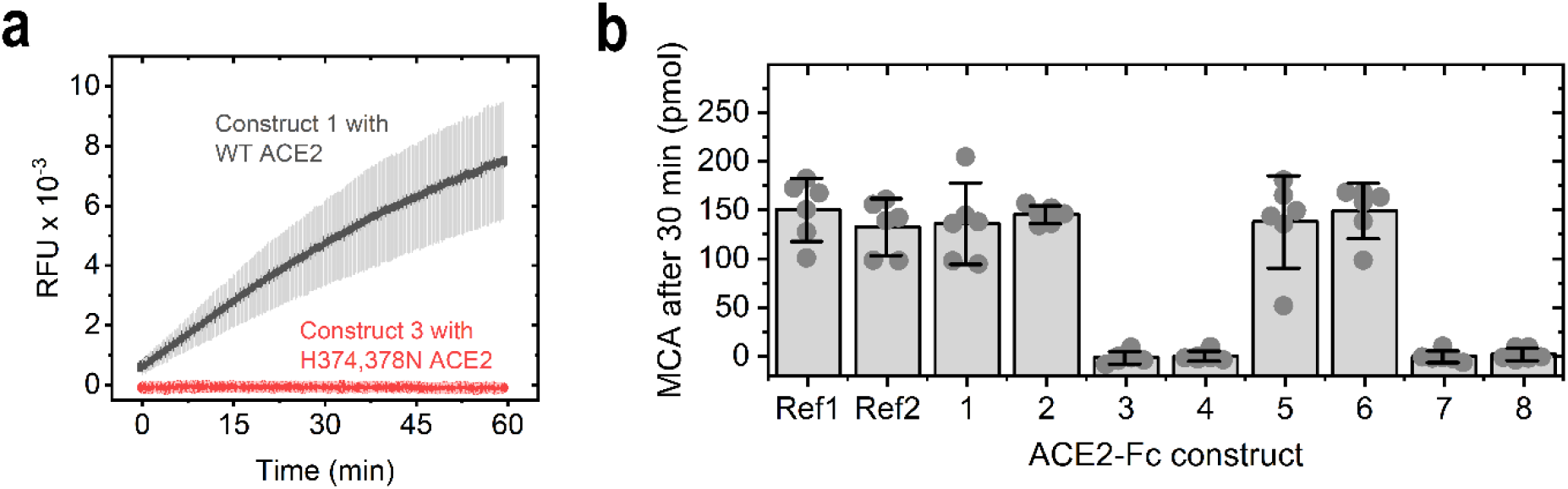
Enzymatic activity of the ACE2-Fc constructs. a. Comparison of fluorescence signals over time obtained in an assay testing the cleavage of a fluorescent peptidyl-4- methylcoumaryl-7-amide (MCA). Relative fluorescent units (RFU) are given. Comparison of the cleavage activity of ACE2-Fc construct 1 (containing wildtype ACE2) or construct 3 (containing H374N and H378N mutations expected to ablate enzymatic function) over time. **b** Amount of MCA cleaved after 30 min of incubation with the ACE2-Fc constructs. Ref1 and Ref2 are two different commercially available ACE2-Fc proteins from Genscript and Acrobiosystems, respectively. Bars are mean values; error bars depict the 95% confidence interval of six independent experiments shown as circles.

### Interaction of the ACE2-Fc constructs with the receptor binding domain (RBD) of SARS-CoV-2

Surface plasmon resonance (SPR) allowed to determine the binding affinity of our ACE2-Fc constructs to the RBD of the spike protein of SARS-CoV-2 that was recombinantly expressed and immobilized. The ACE2-Fc fusion proteins bound in a concentration-dependent manner to the viral protein domain, while an unrelated Fc fusion protein (aflibercept) used as a control showed no interaction with the ligand **(Figure 4a)**. The binding constants revealed that all constructs analyzed bind to the immobilized SARS-CoV-2 RBD with a K_D_ of around 4 nM **(Figure 4b)**. This indicated that the structural variations do not influence the interaction between the ACE2-Fc fusion proteins and the RBD of SARS-CoV-2.

**Figure 4.**
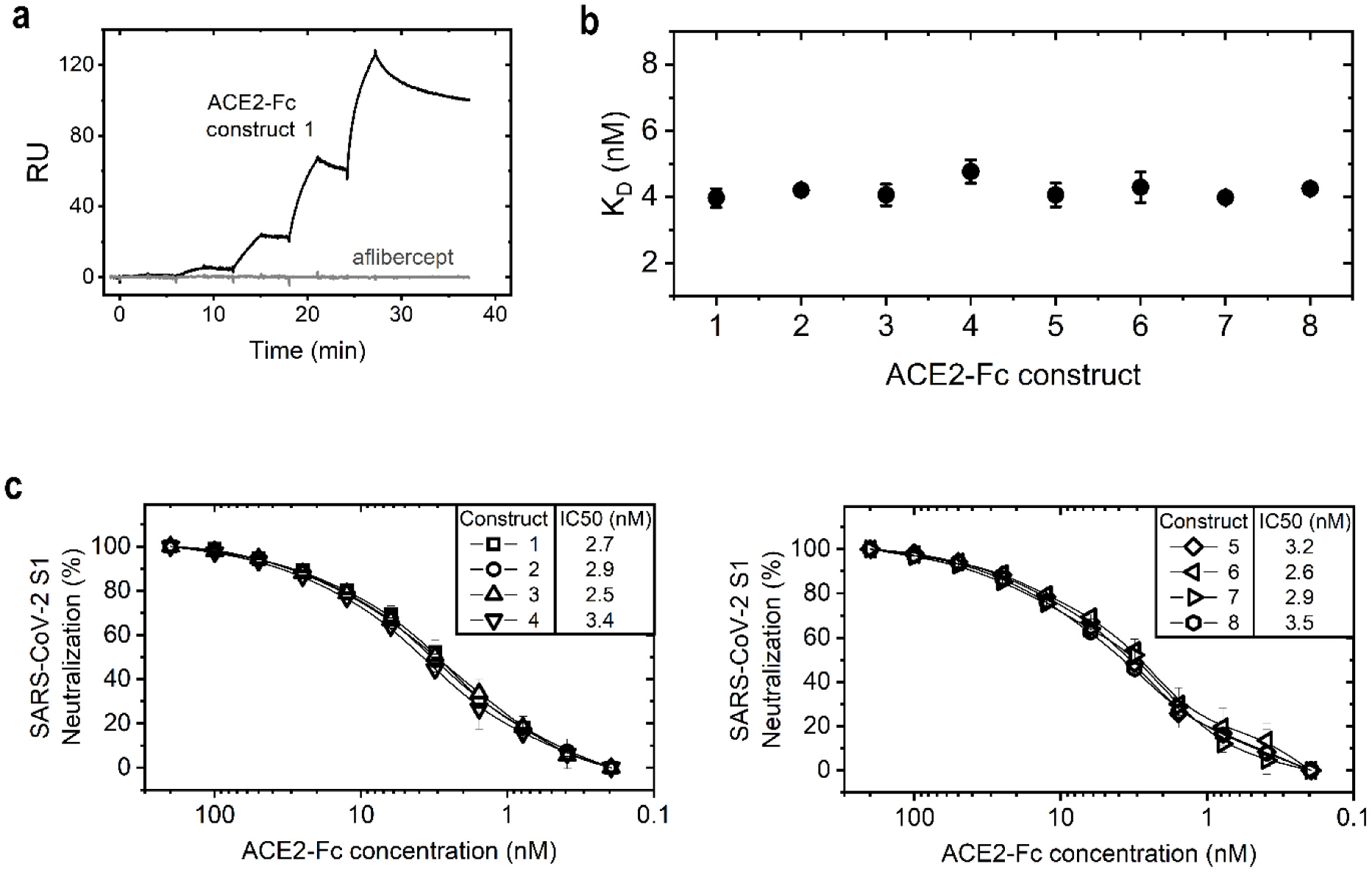
Interaction of the ACE2-Fc constructs with the receptor binding domain. **a** Surface plasmon resonance (SPR) was performed to obtain binding curves of ACE2-Fc fusion constructs and an unrelated Fc fusion protein (aflibercept) to an immobilized RBD from SARS-CoV-2. An exemplary binding curve is shown (RU = Response Units) **b** Binding constants of the ACE2-Fc constructs towards the RBD of SARS-CoV-2 (Mean ±SD of triplicate measurements). **c** ACE2-Fc fusion proteins were pre-incubated with the SARS-CoV-2 spike S1 protein and tested in a competition ELISA for their ability to neutralize S1 binding to immobilized ACE2 protein. Potent inhibition of SARS-CoV-2 spike S1 protein by ACE2-IgG4-Fc constructs (left) and ACE2-IgG1-Fc constructs (right). Data are means ± SD of at least two independent experiments.

Neutralizing activities of the ACE2-Fc fusion proteins against the SARS-CoV-2 spike protein were tested in a competition enzyme-linked immunoassay (ELISA). All ACE2-Fc constructs tested bound and potently neutralized the binding of spike S1 protein of SARS-CoV-2 to ACE2 **(Figure4c)**. Consistent with their affinities to the SARS-CoV-2 RBD, there were no significant differences between the different fusion proteins. The half-maximal inhibitory concentrations (IC50 values) for SARS-CoV-2 spike S1 protein neutralization ranged from 2.5 to 3.5 nM.

### Neutralization assay with ACE2-IgG4-Fc using SARS-CoV-2-GFP

We next wanted to test if the competition of our ACE2-Fc fusion constructs with binding of the SARS-CoV-2 *via* S1 to its receptor ACE2 would translate into neutralization of infectious virus. Because of the expected favorable *in vivo* features, we chose two IgG4-based ACE2-Fc fusion proteins, construct 1 and construct 3, for virus neutralization tested *via* live cell imaging using the IncuCyte platform in a biosafety level 3 laboratory. When Vero E6 cells were infected with SARS-CoV-2-GFP [39] pre-incubated with serial dilutions of the two ACE2-IgG4-Fc fusion proteins (ACE2 Q18-G732 wildtype and ACE2 Q18-G732 H374N/H378N fused to IgG4-Fc S228P) the virus was neutralized in a concentration-dependent manner and no green fluorescent protein (GFP) expression was detected in contrast to cell layers showing increasing GFP expression when infected with non-treated virus. **(Figure 5 and Supplementary Movies)**. This demonstrates that construct 1 and construct 3 completely reduce SARS-CoV-2-GFP infection of Vero E6 cells *in vitro.*

**Figure 5.**
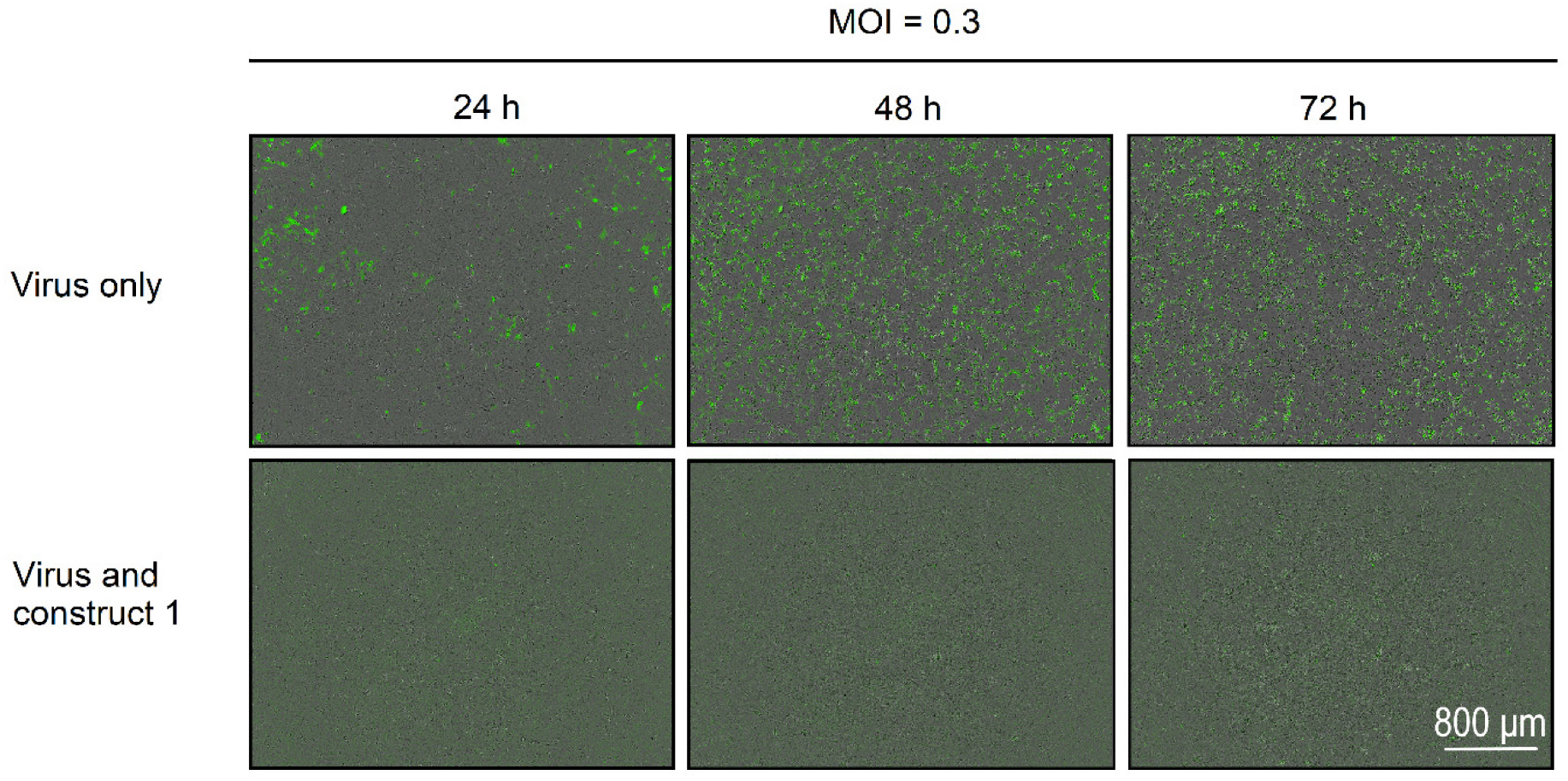
ACE2-IgG4-Fc dependent reduction of SARS-CoV-2-GFP replication. Representative fluorescent images of Vero E6 cells infected with SARS-CoV-2-GFP (Multiplicity of infection (MOI) = 0.3) pre-incubated with ACE2-IgG4-Fc fusion construct 1 (632 nM) (scale bar: 800 μm).

### Neutralization of a variety of SARS-CoV strains

To determine whether the ACE2-Fc constructs would be able to neutralize a broader range of SARS-coronavirus strains, we compared the neutralization capacity of each of the eight ACE2-Fc fusion constructs against a range of virus isolates. Hereby, we included the original SARS-CoV isolate from 2003 as well as two different SARS-CoV-2 strains isolated in January and April 2020. All ACE2-IgG-Fc fusion proteins developed against the novel SARS-CoV-2 also neutralized the first human pathogenic SARS-CoV circulating in 2003 [40] with 50% inhibitory concentration (IC50) values in the range of 150 nM **(Figure 6a)**

**Figure 6.**
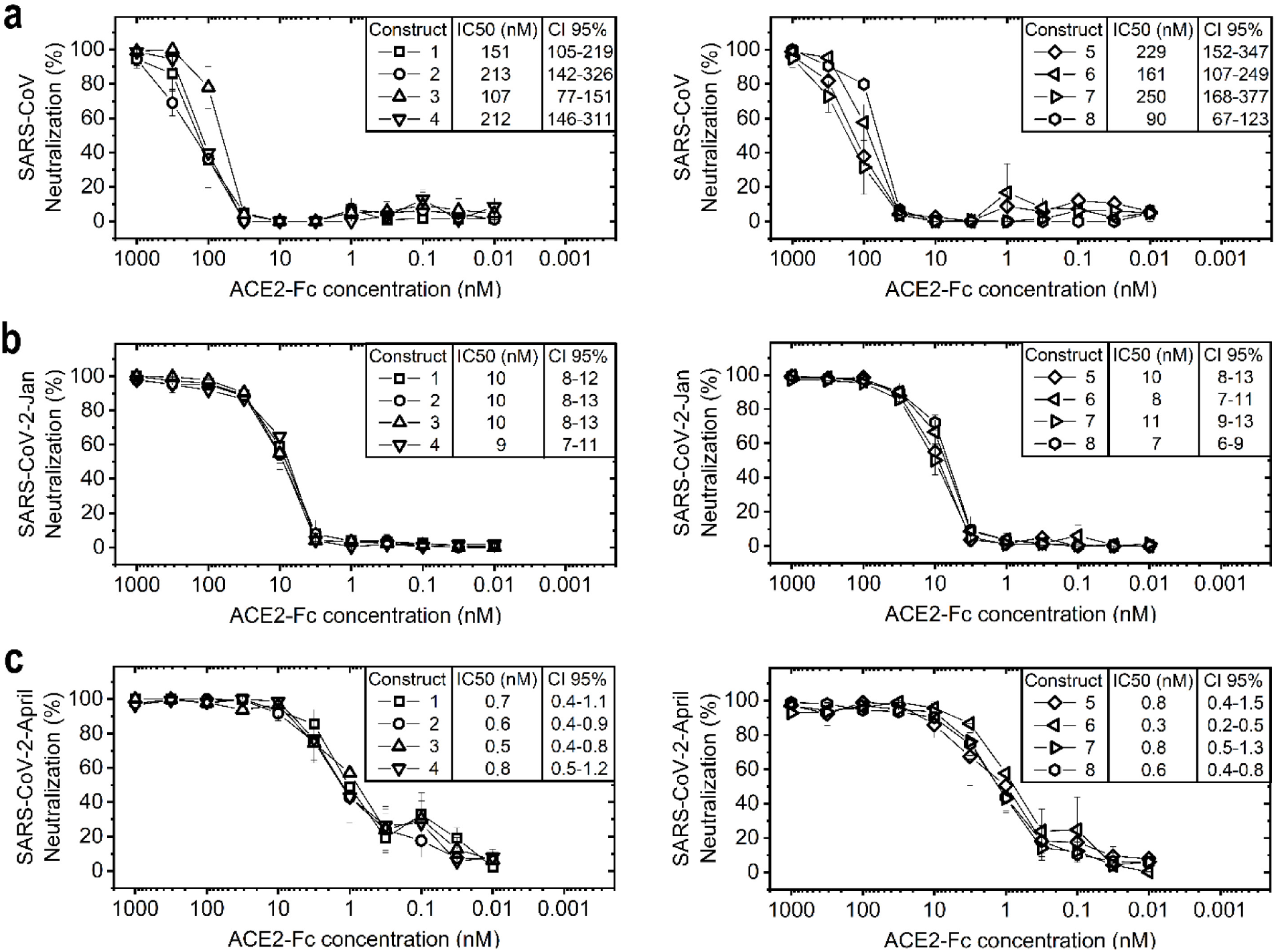
ACE2-Fc fusion proteins potently neutralize coronaviruses. Serial dilutions of ACE2-Fc fusion proteins were pre-incubated with different coronaviruses and tested for their ability to neutralize the virus before infection of Vero E6 cells. Neutralization of SARS-CoV **(top)**, SARS-CoV-2-Jan **(middle)** and SARS-CoV-2-April **(bottom)** by ACE2-IgG4-Fc constructs **(left)** and ACE2-IgG1-Fc constructs **(right)** is shown. Data given are means ± SEM of three independent experiments each. 50% inhibitory concentrations (IC50) determined as well as the 95% confidence interval (CI 95%) are given for each construct.

We then tested for the ability of the ACE2-Fc constructs to neutralize SARS-CoV-2, in particular SARS-CoV-2-Jan isolated from a Bavarian COVID-19 patient [41] from the earliest documented COVID-19 outbreak in Germany end of January 2020 [42], which was directly connected to the initial outbreak in Wuhan, China. All ACE2-Fc fusion proteins displayed strong neutralizing potential against SARS-CoV-2-Jan **(Figure 6b)** with IC50 values in the range of 10 nM.

The ACE2-Fc fusion protein variants were also compared for their ability to neutralize a second SARS-CoV-2 isolate, SARS-CoV-2-April, which was isolated when the virus was massively spreading in Europe. SARS-CoV-2-April displayed a different plaque-forming phenotype than SARS-CoV-2-Jan **(Supplementary Figure 1)**. All ACE2-Fc constructs displayed significantly increased neutralizing potential against SARS-CoV-2-April **(Figure 6c)** with IC50 values below 1 nM.

## Discussion

In this study, we design, express and evaluate different ACE2-Fc fusion proteins with favorable biochemical features and show that they elicit a broad antiviral activity against different human pathogenic coronaviruses including the 2003 SARS-CoV, as well as different strains of the 2020 SARS-CoV-2. All ACE2-IgG4-Fc and ACE2-IgG1-Fc fusion proteins selected inhibited infection of Vero E6 cells by the SARS-CoV-2 wildtype virus isolate SARS-CoV-2-Jan, which was obtained from the earliest documented COVID-19 cases in Germany from January 2020 and thus is closely related to the original Wuhan strain [41, 42], with IC50 values of around 10 nM. Infection by the predominant SARS-CoV-2-April variant circulating worldwide was inhibited even more efficiently with IC50 values below 1 nM. All ACE2-Fc molecules also inhibited infection with the original SARS-CoV with IC50 values in the range of 150 nM. These results demonstrate that the ACE2-Fc constructs can be used for neutralization of various coronavirus strains.

Various sequence variants of ACE2-IgG1-Fc fusion proteins have been described in the literature [21–26]. In this study, novel ACE2-Fc fusion proteins in which the ACE2 domain is fused to the Fc fragment of human IgG 4 (ACE2-IgG4-Fc) containing a stabilizing S228P mutation in the hinge region were designed. We used the full length ACE2 ectodomain sequence Q18-S740 and a shortened version comprising Q18-G732. In addition, an ACE2 variant was used comprising two point mutations which abolish its enzymatic activity.

It was thought that ACE2-Fc fusion proteins are homodimers stabilized by disulfide bonds in the hinge region, similar to IgGs [43]. The ACE2 domain and the Fc part likely fold independently, however the Fc domain might have an impact on the overall stability of the molecule and the interaction with the virus spike protein. We therefore wondered whether the Fc fragment isotype impacts expression, purity, and activity of the ACE2 fusion protein. A comparison of the ACE2-IgG4-Fc and ACE2-IgG1-Fc constructs showed that they express equally well in mammalian cells, suggesting a similar *in vivo* folding and assembly efficiency, although yield and purity was slightly higher for the shortened Q18-G732 ACE2 sequence. Also, the enzymatic activity of the ACE2 domain is not affected by the Fc fusion in the wildtype constructs. As expected, the enzymatic activity was completely abolished in the ACE2 double assay point mutants. Thus, the basic properties of ACE2 do not seem to be affected in the two different fusion formats. In addition, the binding to the SARS-CoV-2 viral spike protein as determined by SPR is similar with a K_D_ of 4 nM. Very similar values were also observed in an ELISA-based SARS-CoV-2 spike S1 protein neutralization assay for all ACE2-IgG4-Fc and ACE2-IgG1-Fc constructs.

Investigations on the genome sequences of multiple SARS-CoV-2 isolates reported from various countries identified genomic regions with increasing genetic variation [44–46]. Globally, one of these variations, the D614G substitution in the C-terminal region of the spike protein, is now the most prevalent pandemic form [47, 48]. Mutation D614G is associated with an improved ability to bind to ACE2 protein and infection. In addition, the D614G substitution makes the spike protein more sensitive for some neutralizing antibodies [49–51]. Our results indicate that both, the ACE2- IgG4-Fc fusion proteins and the ACE2-IGg1-Fc fusion proteins, even more potently neutralize the worldwide predominant SARS-CoV-2-April variant than SARS-CoV-2-Jan which is closely related to the non-mutated, original SARS-CoV-2 strain from Wuhan. These findings are consistent with previous work that has shown that the D614G mutation increases the susceptibility of the virus towards neutralization [50, 51]. In line with our results, the affinity of the SARS-CoV-2 spike protein for human ACE2 has been previously reported to be significantly higher than the affinity of the SARS-CoV spike protein [3]. This may explain the high replication rate of SARS-CoV-2 in the upper respiratory tract and thus the high rate of infection.

Here we describe a decrease of the IC50 in the neutralization assays from 10 nM to 1 nM with SARS-CoV-2 isolates obtained from patients in Germany in January 2020 and April 2020 which indicates a further affinity maturation of the SARS-CoV-2 spike protein towards human ACE2 during the course of the pandemic.

It remains a concern that SARS-CoV-2 and other coronavirus variants have found reservoirs in animals living in close contact with humans and could transmit to humans from time to time, as happened recently with “cluster 5”, a SARS-CoV-2 variation with a combination of mutations that is not fully understood for its impact on disease severity and resistance to vaccination and therapeutic antibodies [52, 53]. In contrast to vaccines and therapeutic antibodies, which have been associated with a risk of virus escape by mutation as e.g. the occurrence of the spike variant N439K [54–57], administration of ACE2 as a virus blocker provides a high level of protection against drug resistance caused by virus mutation.

ACE2 plays a central role in the homeostatic control of cardio-renal actions and has been shown to protect against severe acute lung injury and acute angiotensin II-induced hypertension [58, 59].

Moreover, ACE2 has been identified as a functional receptor for SARS-CoV, SARS-CoV-2 and NL63 [1–3] which is confirmed by our data showing neutralization potency for both SARS-CoV and SARS-CoV-2. Recombinant human ACE2, however, also is an interesting therapeutic candidate for COVID-19 [20, 27]. A soluble dimer (APN019) has been safely tested in a clinical phase I study in healthy volunteers and in a phase II study in patients with an acute respiratory distress syndrome [18, 19]. It is currently being tested for its therapeutic effect in a phase II study in COVID-19 patients [20]. A human ACE2-Fc fusion protein designed with an IgG1 Fc portion to prolong the circulating half-life is currently in a phase I clinical study pursued by Hengenix Biotech Inc. [27].

Pharmacokinetic studies in mice and humans revealed that recombinant human ACE2 exhibits fast clearance rates resulting in a short half-life of only a few hours [18, 20, 58]. When the extracellular domain of murine ACE2 was fused to an Fc fragment of murine immunoglobulin IgG the ACE2-Fc fusion protein demonstrated a prolonged *in vivo* half-life and effective organ protection in murine models of both, acute and chronic angiotensin II-dependent hypertension [60].

The presence of the Fc domain markedly increases the plasma half-life of ACE2-Fc due to the interaction of the Fc domain with the neonatal Fc-receptor (FcRn), and therefore a slower renal clearance of the fusion molecule. FcRn is broadly expressed on many cell types including endothelial cells and respiratory epithelial cells [61, 62]. Binding to FcRn extends the systemic half-life by chaperoning bound Fc fusion proteins away from lysosomal degradation. In addition, FcRn transports IgG and Fc fusion molecules across mucosal barriers into the lumen of the respiratory and intestinal tract thereby providing a dynamic trafficking between circulating and luminal IgG molecules at mucosal sites [61, 63].

The affinity of ACE2 dimer to SARS-CoV-2 spike protein has been reported to be lower compared to an ACE2-IgG1-Fc fusion protein [22, 26]. Using our design, ACE2-IgG4-Fc and ACE2-IgG1-Fc have both shown high binding affinity to the SARS-CoV-2 RBD and spike protein and bound and neutralized the virus with IC50 values below 1 nM. In addition, the IgG4-Fc fragment could reduce the risk of disease enhancement related to complement dependent cytotoxicity (CDC), antibody dependent cytotoxicity (ADCC) and infection *via* CD16 Fc receptor [31]. Although the constant heavy chain regions of different IgG subclasses share over 95% sequence homology, their structures and effector functions differ. IgG4 in particular has poor ability to engage C1q and Fc gamma receptors and has been associated with anti-inflammatory properties [64].

The SARS-CoV-2 pandemic has caused an unprecedented challenge to develop COVID-19 drugs in rapid pace. However, progress with candidate therapies still has been slow. Here we show that ACE2-IgG4-Fc fusion proteins have favorable biophysical and pharmaceutical characteristics and significant *in vitro* SARS-CoV-2 neutralizing potency. The Fc part from IgG4 could bring clinical benefits for ACE2-Fc proteins by avoiding potential antibody-dependent disease enhancement during treatment. Therefore, ACE2-IgG4-Fc fusion proteins present a promising potential treatment option not only for the current SARS-CoV-2 pandemic but also for future coronavirus infectious diseases.

## Materials and Methods

### Construct design

ACE2 amino acid sequence modifications were designed by computer-aided modelling. ACE2 ectodomains of different length, Q18-G732 and Q18-S740, with or without mutation of the catalytic site (wild type or H374N/H378N mutant) [65] were combined with the Fc fragment of IgG4 bearing a stabilizing S228P mutation in the hinge region [33]. For comparison, the same ACE2 sequence variants were fused to the Fc fragment of IgG1 with a truncated hinge region (DKTHTCPPCPA).

### Generation of expression plasmids

Plasmids encoding the Fc fusion proteins were generated at ThermoFisher. Genes of interest were subcloned into pcDNA3.1 Zeocin expression plasmids (Invitrogen V860-20) with an elongated CMV promoter using HindIII/XhoI restriction sites. Following amplification in *Escherichia coli*, expression plasmids were isolated and analyzed by restriction analysis as well as DNA sequencing.

### Protein expression

Using the FreeStyle 293 Expression System (ThermoFisher), the different ACE2-Fc fusion proteins were transiently expressed in 3 x 240 mL culture media. On day six, samples were analyzed for cell viability as well as cell density and supernatants were harvested by centrifugation followed by sterile-filtration [66]. The material was either stored at −80°C until purification or subjected directly to purification. Small samples were taken from the pools to determine expression yields by bio-layer interferometry (BLI).

### Protein purification

Purification of the fusion proteins secreted into the culture medium was performed by protein A column chromatography followed by preparative Size Exclusion Chromatography (SEC). For protein A purification, after loading the sample, the Amsphere A3 column (JSR Life Sciences) was washed and the ACE2-Fc fusion proteins were eluted using 40 mM NaAc, pH 3.0. Following elution, samples were first neutralized to pH 7.5 using 1 M Tris, pH 9.0, subsequently diluted 1:1 with 50 mM Tris, pH 7.5, 300 mM NaCl and concentrated to 10 mg/mL using spin filters. Concentrated proteins were further purified with a Superdex 200 increase (GE Healthcare) column equilibrated with 50 mM Tris, pH 7.5, 150 mM NaCl. The main peak was pooled, the protein concentration was determined by slope spectrometry [67] and adjusted to 1 mg/mL. The protein solution was passed through a sterilizing filter and stored at 4°C until further usage.

### Size exclusion chromatography with multi-angle light scattering (SEC-MALS)

A Shimadzu HPLC system with two concentration detectors (UV and refractive index) and a HELEOS II MALS detector were used for the measurements. The flow rate was 1 mL/min and the running buffer was 50 mM Tris, pH 7.5 and 150 mM NaCl. 50 μg of protein was injected on a Superdex 200 Increase 10/300 GL column (Cytiva). The chromatograms were evaluated with the Astra software.

### Circular dichroism

All CD measurements were performed with a Jasco J-1500 spectropolarimeter at 20°C. The sample buffer consisted of 50 mM Tris, pH 7.5 and 150 mM NaCl. The Far-UV CD spectra were obtained in a 1 mm quartz cuvette using a protein concentration of 0.1 mg/mL. The Near-UV CD spectra were measured in a 5 mm quartz cuvette using a protein concentration of 1 mg/mL.

### ACE2 activity assay

An ACE2 activity assay kit from Abcam (Cat.No. ab273297) was used to measure the enzymatic activity of the constructs. The assay was performed according to the manufacturer’s manual and is based on a synthetic peptidyl-4-methylcoumaryl-7-amide (MCA) that is cleaved by the ACE2 enzyme. Upon cleavage, free MCA is detected fluorometrically (Ex320nm/Em420nm) and quantified with a standard curve obtained with MCA standard solutions with known concentrations. Two commercially available ACE2-Fc proteins obtained from Genscript (Cat.No. Z03484-1) and Acrobiosystems (Cat.No. AC2-H5257) were used as reference.

### Determination of binding affinity to spike protein RBD using surface plasmon resonance

The measurements were performed with a Biacore X-100 system and the Biotin CAPture kit (Cytiva). The running buffer was HBS-EB+ (Cytiva). The ligand, SARS-CoV-2 RBD with an AviTag (Acrobiosystems) was immobilized on the streptavidin chip to around 100 RU. Increasing concentrations of the analyte ACE2-Fc (0.32, 1.6, 8, 40 and 200 nM) were injected over the immobilized ligand in a single-cycle kinetic mode. The obtained sensorgrams were evaluated with the Biacore X-100 software to obtain a binding constant (KD).

### SARS-CoV-2 Spike S1 Inhibition ELISA

Inhibition of binding of SARS-CoV-2 spike S1 protein to ACE2 was tested using the ACE2:SARS-CoV-2 Spike S1 Inhibitor Screening Assay Kit (BPS Bioscience; Cat.No. 79945) according to the manufacturer’s instructions with an adapted neutralization procedure. Briefly, biotinylated SARS-CoV-2 Spike S1 protein (25 nM) was incubated with serial dilutions of the ACE2-Fc fusion proteins in a 96-well neutralization plate at room temperature (RT) for one hour with slow shaking (= neutralization mix).

ACE2 protein was added to a nickel-coated 96-well plate at a concentration of 1 μg/mL and incubated at RT for one hour with slow shaking. Following a washing step to remove unbound ACE2, the plates were blocked at RT for 10 min with slow shaking. Subsequently, the neutralization mix was transferred to the ACE2 coated plate and the plate was incubated at RT for one hour with slow shaking. Following a 10 min blocking step, the plate was incubated with Streptavidin-HRP at RT for one hour with slow shaking. Following a washing and a 10 min blocking step, the HRP substrate was added and the plate was analyzed on a chemiluminescence reader.

### Virus strains

SARS-CoV-2-GFP (kindly provided by Volker Thiel, University of Bern, Switzerland) is based on the original Wuhan SARS-CoV-2 isolate (GenBank accession MT108784) and was reconstituted from a synthetic construct derived from SARS-2 BetaCoV/Wuhan/IVDC-HB-01/2019 [39].

SARS-CoV-2-Jan (SARS-CoV-2-Munich-TUM-1; EPI_ISL_582134), SARS-CoV-2-April (SARS-CoV-2 D614G; EPI_ISL_466888), and SARS-CoV (AY291315.1) were isolated from patient material in Germany. Briefly, SARS-CoV-2-Jan was isolated from a COVID-19 infected patient who was infected during the earliest documented COVID-19 outbreak in Germany at the end of January 2020 with a virus imported from Wuhan *via* a single contact in Shanghai [41]. The SARS-CoV-2-April strain was isolated during the first eminent wave of the pandemic in Europe in April 2020 from a patient in Munich. Both virus isolates as well as a control isolate from the early “Webasto” cluster outbreak [68] contain the S1 D614G mutation showing significantly higher infectious titers *in vitro*[47].

SARS-CoV-2-Jan, SARS-CoV and SARS-CoV-2-GFP [39] were propagated and passaged in Vero E6 cells (derived from African green monkey kidney epithelial cells). SARS-CoV-2-April was propagated by the infection of Caco-2 cells followed by passaging in Vero E6 cells. All strains were cultured in DMEM medium (5% fetal calf serum (FCS), 1% penicillin/streptomycin (P/S), 200 mmol/L L-glutamine, 1% MEM-Non-Essential Amino Acids (NEAA), 1% sodium-pyruvate (all from Gibco). Viral titer was determined by Plaque Assay [69].

### Plaque Assay

Viral titers were determined as described by Baer et al., [69] with some modifications. Briefly, HepG2 or Vero E6 cells were plated in a 12-well plate at 5 x 10E5 cells/well in DMEM medium (Gibco) supplemented with 5% FCS, 1% P/S, 200 mmol/L L-glutamine, 1% MEM-NEAA, 1% sodium-pyruvate (all from Gibco) and incubated overnight at 37°C and 5% CO2. Cells were infected with serial dilution of virus sample in cell culture medium at 37°C for one hour. After discarding the supernatant, 1 mL of 5% carboxymethylcellulose (Sigma) diluted in Minimum Essential Media (Gibco) was added per well and the plate was incubated at 37°C until obvious plaques appeared. After removing the supernatant, cells were fixed with 10% paraformaldehyde (ChemCruz) at RT for 30 min. Next, a washing step with PBS was performed, followed by the addition of 1% crystal violet (Sigma; diluted in 20% methanol and water). Following an incubation time of 15 min at RT, the solution was washed away with PBS and the plate was dried. The viral titer (PFU/mL) of the sample was determined by counting the average number of plaques for a dilution and the inverse of the total dilution factor.

### SARS-CoV / SARS-CoV-2 virus neutralization assay

#### Viral neutralization assay using SARS-CoV-2-GFP

Vero E6 cells were plated in a 96-well plate at 1.4 x 10E04 cells/well in DMEM medium (Gibco) supplemented with 5% FCS, 1% P/S, 200 mmol/L L-glutamine, 1% MEM-NEAA, 1% sodium-pyruvate (all from Gibco) and incubated overnight at 37°C and 5% CO2. Serial dilutions of ACE2-Fc fusion proteins and SARS-CoV-2-GFP were mixed in fresh media and pre-incubated at 37°C for one hour. Afterwards, Vero E6 cells were infected at a multiplicity of infection (MOI) of 0.3 infectious viruses per cell at 37°C. After one hour, the neutralization mix was replaced by cell culture medium. Plates were placed in the IncuCyte S3 Live-Cell Analysis System and real-time images of uninfected mock cells (Phase channel) and infected (GFP and Phase channel) cells were captured every four hours for 72 hours. Virus control cells were infected with the same virus stock but without prior incubation with the ACE2-Fc fusion constructs using the identical protocol.

#### Viral neutralization assay followed by in-cell ELISA

Vero E6 cells were plated in a 96-well plate at 1.6 x 10E04 cells/well in DMEM medium (Gibco) supplemented with 5% FCS, 1% P/S, 200 mmol/L L-glumatine, 1% MEM-NEAA, 1% sodium-pyruvate (all from Gibco) and incubated overnight at 37°C and 5% CO2. Serial dilutions of the ACE2-Fc fusion proteins were mixed with virus in fresh media and pre-incubated at 37°C for one hour. The Vero E6 cells were infected at a multiplicity of infection (MOI) of 0.03 with the neutralized virus solution at 37°C for one hour. Next, the neutralization mix was removed, culture medium was added, and cells were incubated at 37°C for 24 hours. Mock cells represent uninfected Vero E6 cells, incubated with culture medium. After 24 hours, cells were washed once with PBS and fixed with 4% paraformaldehyde (ChemCruz) at RT for 10 min. Following a washing step with PBS, fixed Vero E6 cells were permeabilized with 0.5% saponin (Roth) in PBS at RT for 10 min. Next, the permeabilization solution was removed and cells were blocked with a mixture of 0.1% saponin and 10% goat serum (Sigma) in PBS with gentle shaking at RT for one hour. Subsequently, Vero E6 cells were incubated with a 1:500 dilution of an anti-dsRNA J2 antibody (Jena Bioscience) in PBS supplemented with 1% FCS at 4°C overnight with shaking. Following four washing steps with wash buffer (PBS supplemented with 0.05% Tween-20 (Roth)). Next, the plates were incubated with a 1:2,000 dilution of a goat anti-mouse IgG2a-HRP antibody (Southern Biotech) in PBS supplemented with 1% FCS and incubated with gently shaking at RT for one hour. Following four washing steps, 3,3’,5,5’-Tetramethylbenzidin (TMB) substrate (Invitrogen) was added to the wells and incubated in the dark for 10 min. Colorimetric detection on a Tecan infinite F200 pro plate reader at 450 nm and at 560 nm was performed after stopping the color reaction by the addition of 2N H2SO4 (Roth).

## Supporting information

Constrcut 1 plus SARS-CoV-2-GFP

Medium Only

SARS-CoV-2-GFP Only

## Data Availability

The authors declare that the data supporting the findings of this study are available within the article and its Supplementary Information files, or are available from the authors upon request. Source data are provided with this paper.

## Acknowledgements

The authors like to thank Polpharma Biologics Utrecht B.V. for performing the transient transfections and providing the fusion molecules. We are grateful to Volker Thiel, University of Bern, Switzerland, for providing the SARS-CoV-2-GFP, and to Friedemann Weber, University of Giessen, Germany, for providing SARS-CoV (Frankfurth1).

Supported by grant AZ-1433-20 of the Bayerische Forschungsstiftung.

## Supplementary Material and Methods

### Analytical Size Exclusion Chromatography (SEC)

Purified proteins were analyzed by analytical size exclusion chromatography (SEC) on a Waters H-Class bio UPLC system using an Acquity UPLC Protein BEH SEC column, 4.6 mm x 150 mm, 200 Å, 1.7 μm. Detection was based on UV absorbance at 280 nm. 20 μg of samples were loaded, the mobile phase consisted of 20 mM sodium phosphate pH 7.0, 150 mM NaCl and proteins were eluted isocratically at a flow rate of 0.3 mL/min.

### Capillary Electrophoresis - Sodium Dodecyl Sulfate (CE-SDS)

Purified proteins were analyzed by capillary electrophoresis sodium dodecyl sulfate (CE-SDS) under non-reducing as well as reducing conditions on a CESI8000 instrument (AB Sciex) following the manufacturer’s protocol from the IgG purity/heterogeneity kit. Before the measurement *via* pressure injection, the buffer was adjusted to 50 mM Tris pH 7.5, 50 mM NaCl.

### Supplementary Figure

**Supplementary Figure 1.**
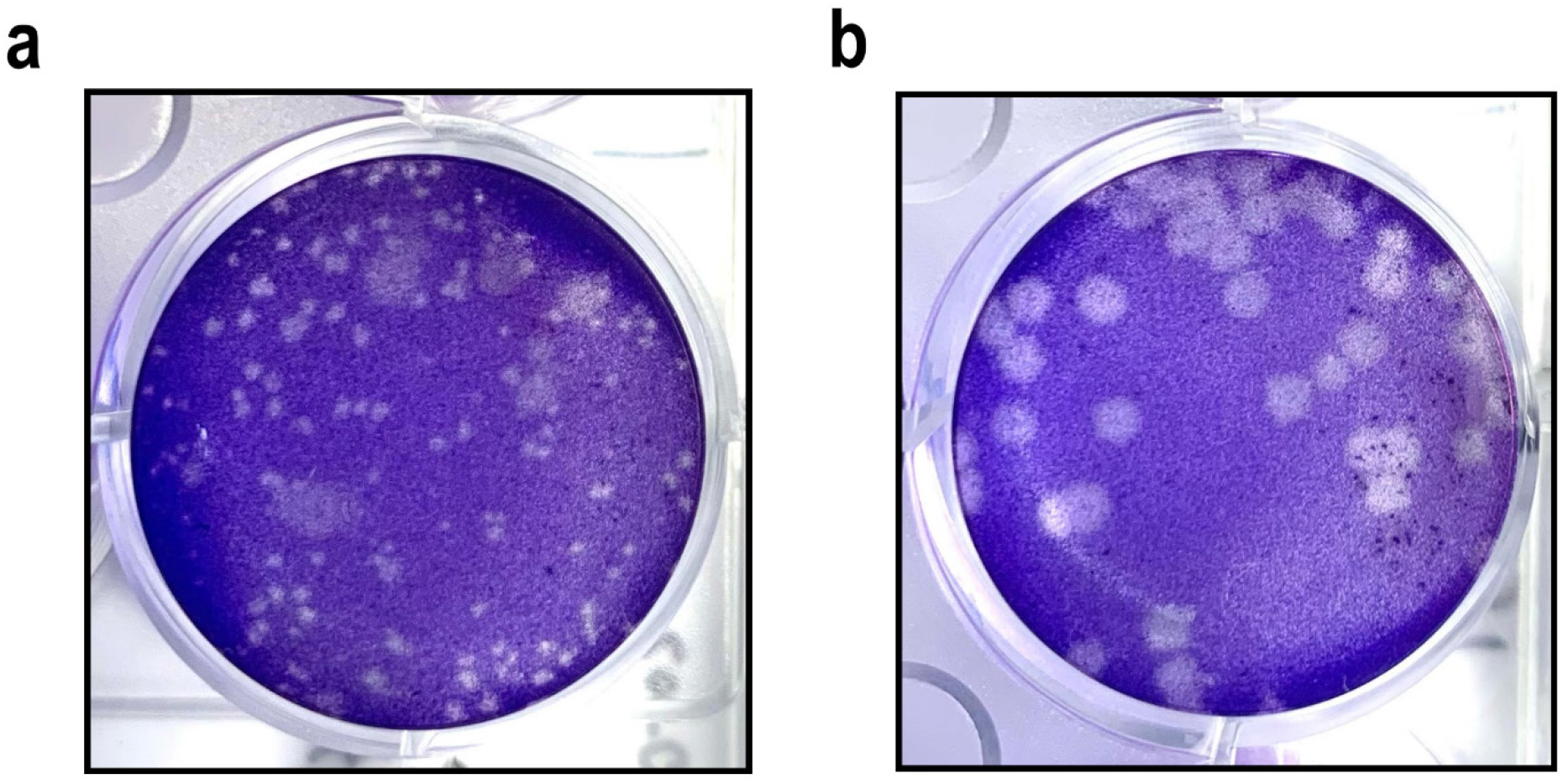
Plaque appearance. Viral plaques following infection of HepG2 cells with **a** SARS-CoV-2-Jan or **b** SARS-CoV-2-April.

### Supplementary Table

**Supplementary Table 1.** Yield and purity of the ACE2-Fc constructs

| **Construct** | **Normalized**<br>**Relative Yield\*** | **HMWS (%)**<br>**Analytical SEC** | **Purity**<br>**CE-SDS non-reduced** | **Purity**<br>**CE-SDS reduced** |
| --- | --- | --- | --- | --- |
| **1** | 0.8 | 2.1 | 97.3 | 96.7 |
| **2** | 0.6 | 4.7 | 97.4 | 97.4 |
| **3** | 1.0 | 3.1 | 95.1 | 94.8 |
| **4** | 0.6 | 3.7 | 97.0 | 98.2 |
| **5** | 1.0 | 2.6 | 97.0 | 95.2 |
| **6** | 0.6 | 3.4 | 97.2 | 96.8 |
| **7** | 0.9 | 1.9 | 95.3 | 96.7 |
| **8** | 0.8 | 3.7 | 96.9 | 97.8 |

\* Normalized against construct 5

